# Transposable element diversity remains high in gigantic genomes

**DOI:** 10.1101/2021.08.27.457980

**Authors:** Ava Louise Haley, Rachel Lockridge Mueller

## Abstract

Transposable elements (TEs) are repetitive sequences of DNA that replicate and proliferate throughout genomes. Taken together, all the TEs in a genome form a diverse community of sequences, which can be studied to draw conclusions about genome evolution. TE diversity can be measured using models for ecological community diversity that consider species richness and evenness. Several models predict TE diversity decreasing as genomes expand because of selection against ectopic recombination and/or competition among TEs to garner host replicative machinery and evade host silencing mechanisms. Salamanders have some of the largest vertebrate genomes and highest TE loads. Salamanders of the genus *Plethodon*, in particular, have genomes that range in size from 20 to 70 Gb. Here, we use Oxford Nanopore sequencing to generate low-coverage genomic sequences for four species of *Plethodon* that encompass two independent genome expansion events, one in the eastern clade (*P. cinereus*, 29.3 Gb vs. *P. glutinosus*, 38.9 Gb) and one in the western clade (*P. vehiculum*, 46.4 Gb vs *P. idahoensis*, 67.0 Gb). We classified the TEs in these genomes and found ~52 TE superfamilies, accounting for 27-32% of the genomes. We calculated Simpson’s and Shannon’s diversity indices to quantify overall TE diversity. In both pairwise comparisons, the diversity index values for the smaller and larger genome were almost identical. This result indicates that, when genomes reach extremely large sizes, they maintain high levels of TE diversity at the superfamily level, in contrast to predictions made by previous studies on smaller genomes.

## INTRODUCTION

Genome sizes vary ~300,000-fold among eukaryotes, from ~ 0.002 Gb (e.g. in the eukaryotic fungi *Encephalitozoon cuniculi*) to ~670 Gb (e.g. in *Amoeba dubia*). Across animals, the differences span 6,650-fold (Gregory 2021). Salamanders, one of the three clades of amphibians comprising 765 extant species (AmphibiaWeb 2021), include many of the largest animal genomes, ranging from ~9 Gb in *Thorius spilogaster* to 120 Gb in *Necturus lewisi* (Decena-Segarra, et al. 2020; Gregory 2021). The main proximate cause for their large and variably sized genomes is the proliferation of transposable elements (Sun, Shepard, et al. 2012), which contribute to differences in genome size across diverse taxa (Wells and Feschotte 2020). Transposable elements (TEs) are DNA sequences that replicate and insert themselves throughout the genome. The percentage of the genome made up of TEs varies greatly across the tree of life, from ~ 0.1% (e.g. in the fungi *Pseudozyma antarctica*) to ~90% (e.g. in the lily *Fritillaria imperialis*) (Ambrozova, et al. 2011; Castanera, et al. 2017). In salamander genomes, ~25% - ~50% of the total DNA has been classified as recognizable TEs depending on the species (Sun, Shepard, et al. 2012; Sun and Mueller 2014; Nowoshilow, et al. 2018). Because the majority of TEs serve no initial protein-coding or regulatory function in the genome, they accumulate mutations, causing them to be undetectable during TE annotation (Venner, et al. 2009). Thus, the percentages in salamanders do not include older TE insertions that have accumulated mutations and become unrecognizable, so the genomes are likely made up of more TE-derived sequences (Keinath, et al. 2015).

Transposable elements are categorized into two classes. The first is the retrotransposons, which replicate by utilizing the host’s transcriptional machinery to create an RNA intermediate. The RNA intermediate is then reverse-transcribed into a cDNA copy and inserted back into the genome using TE enzymatic machinery (Bourque, et al. 2018). The second is the DNA transposons, which do not have an RNA intermediate and instead move as the direct, excised DNA sequence itself, reinserting into a different location in the genome (Muñoz-López and García-Pérez 2010). Within these classes, TEs are further categorized into 9 orders and > 39 superfamilies, commonly classified using the Wicker unified system (Wicker, et al. 2007), although other classifications also exist (Jurka, et al. 2005; Arkhipova 2017). Many superfamilies can be found in almost all eukaryotes, such as *Gypsy* and *Copia* of the LTR order (Bourque, et al. 2018). Most superfamilies are variable across different genomes, existing at higher or lower proportions depending on the species. For example, in the caecilian *I. bannanicus*, the Class 1 retrotransposon DIRS makes up ~30% of the genome and the retrotransposon LTR/Gypsy makes up ~1%, while in salamanders, LTR/Gypsy is the most abundant (Wang, et al. 2021). In contrast, class II DNA transposons make up 39% - 60% of some teleost genomes, while retrotransposons exist at lower levels (Sotero-Caio, et al. 2017).

Taken together, all of the TEs in a genome form a community of sequences, which can be studied to draw conclusions about genome evolution. As genomes expand, the number of TEs typically increases (Kidwell 2002; Elliott and Gregory 2015b). However, how the diversity of the overall TE community changes with expansion is not yet well understood (Elliott and Gregory 2015a). TE diversity within genomes can be measured in an analogous way to species diversity in ecological communities (Abrusán and Krambeck 2006; Venner, et al. 2009; Linquist, et al. 2015). Analyses of ecological diversity quantify the number of species, or richness, and the abundance of each species, or evenness using the Simpson and Shannon diversity indices (Shannon 1948; Simpson 1949). TE diversity can be approached in a similar way using richness and evenness of TE types (e.g. superfamilies) in a genome (Wang, et al. 2021).

Several analyses have suggested that TE diversity will be highest in smaller genomes. TEs can have negative effects on the fitness of their “hosts” by causing recombination at ectopic, or non-homologous, sites, which can lead to deletions and duplications (Langley, et al. 1988; Petrov, et al. 2003). Because ectopic recombination is more likely to delete or duplicate a functional sequence in smaller genomes, small genome size should select for more diverse TE communities, lowering the number of identical off-target sites to drive errors in crossing-over. In large genomes, the chances of interrupting a functioning gene during ectopic recombination-mediated deletion or duplication are lower. In addition, recombination rates per base pair can be lower, depending on chromosome number, which decreases the likelihood of ectopic recombination overall. Thus, larger genomes can be more permissive to low-diversity TE communities. For these same reasons, larger genomes can be more permissive to TE activity overall, producing a genomic environment in which competition to exploit host replicative machinery, and/or evade host silencing machinery, can lead to a decrease in diversity (Furano, et al. 2004; Abrusán and Krambeck 2006; Boissinot and Sookdeo 2016).

In this study, we test the hypothesis that TE diversity decreases with genome expansion. We chose the salamander genus *Plethodon* (family Plethodontidae) as a study system due to the wide range of genome sizes, but high similarity in physical traits and life history, that exists across the 55 species (Petranka 1998; Gregory 2021). We analyzed two species’ genomes from each of the two main *Plethodon* clades — *P. cinereus* (29.3 Gb genome) and *P. glutinosus* (38.9 Gb) from the eastern clade and *P. vehiculum* (46.4 Gb) and *P. idahoensis* (67.0 Gb) from the western clade, encompassing two independent genome expansion events (Newman, et al. 2016). We rely exclusively on Oxford Nanopore long read sequencing data with no existing genome assembly to reference, demonstrating the power of this method for quantifying TE community diversity. Using both Simpson and Shannon’s diversity indices, we find that TE diversity at the superfamily level remains high as genome size expands. We discuss our findings in light of hypotheses for TE proliferation and silencing dynamics in large genomes.

## MATERIALS AND METHODS

### Tissue Collection

*Plethodon cinereus* and *Plethodon glutinosus* were collected from South Cherry Valley and Oneonta, Otsego County, New York, under the New York State Department of Environmental Conservation scientific collection permit #2303. *Plethodon vehiculum* was collected from Pacific County, Washington, under the Washington Department of Fish and Wildlife scientific collection permit # ITGEN 17-309. *Plethodon idahoensis* was collected in Shoshone County, Idaho, under the Idaho Department of Fish and Game wildlife collection permit #180226. Published genome sizes exist for all four species of *Plethodon* and vary across studies (Gregory 2021), but we use our own lab’s measurements because they were performed on individuals collected at the same time and from the same locality as those sequenced here. Genome sizes for the species are: *P. cinereus* (29.3 Gb), *P. glutinosus* (38.9 Gb), *P. vehiculum* (46.4 Gb), and *P. idahoensis* (67.0 Gb) (Itgen et al, bioRxiv). Animals were euthanized via submersion in 10% buffered MS-222. Tissues were collected and stored in RNALater at -20°C. All work was completed according to the Colorado State University IUCAC protocol (17-7189A).

### DNA Extraction, Library Preparation, and DNA Sequencing

DNA extraction was performed from 0.2 g of trunk skin and muscle tissue using a Qiagen DNEasy Blood and Tissue kit for each species. The manufacturer’s protocol was followed except that 1) samples were flicked instead of vortexed to retain the longest DNA fragments possible, 2) centrifuge times were doubled to ensure all solution passed through the spin column, and 3) 30 μl of elution buffer was used to final increase DNA concentration.

Library preparation was done using a Ligation Sequencing Kit (SQK-LSK109), a Flow Cell Priming Kit (EXP-FLP002), and a Native Barcoding Expansion Kit 13-24 (EXP-NBD114) from Oxford Nanopore. New England Biolabs consumables used were an NEB Blunt/TA Ligase Master Mix (M0367), NEBNext^®^ Quick Ligation Reaction Buffer (NEB B6058), and NEBNext^®^ Companion Module for Oxford Nanopore Technologies^®^ Ligation Sequencing (E7180S). For DNA repair and end prep, the amount of input genomic DNA was increased to 2 μg from the suggested 1 μg. For native barcode ligation, 1000 ng of end-prepped sample was used, twice the amount of suggested sample per the manufacturer’s protocol. A distinct barcode was used for each species. Following barcoding, *P. glutinosus* and *P. cinereus* were pooled together, and *P. vehiculum* and *P. idahoensis* were pooled together to equal about 850 ng of total DNA per pooled sample pair, slightly more than the 700 ng suggested by the protocol. The Long Fragment Buffer was used during adapter ligation. Throughout the protocol, samples were quantified with 1 μl on the Qubit fluorometer. Priming and loading the SpotON flow cell was performed two separate times, with two species occupying one flow cell. Sequencing was performed on the Oxford Nanopore MinION sequencer with the MinKnow software. The sequencer was run for 72 hours with the base calling setting of extremely fast. Porechop was used to trim adapters and barcodes (Wick, et al. 2017).

### Transposable Element Annotation

Our goals were 1) to find the most effective TE annotation tools for low-coverage MinION data possible, enabling accurate calculation of the diversity indices for each genome, and 2) to achieve consistent annotation levels across species, allowing them to be compared without the introduction of bias. In a previous study annotating TEs in the caecilian *Ichthyophis bannanicus*, RepeatMasker and DnaPipeTE together annotated 94.1% of the TE sequences (Wang, et al. 2021). Additionally, in a TE annotation study on the beetle *Dichotomius* (*Luederwaldtinia*) *schiffleriso*, RepeatMasker and DnaPipeTE together annotated 95% of all of the detected TEs in the genome (Amorim, et al. 2020). Although neither study relied on low-coverage MinION data, we chose these two programs together based on these previous successful applications. DnaPipeTE detects TE sequences based on repetitiveness by using Trinity to assemble repeats from low-coverage data. RepeatMasker uses a user-specified library to identify TEs based on sequence similarity. Typically, RepeatMasker is used to mask detected TEs from the genome of interest in order to allow analysis of the non-repetitive portions, but for studies focused on TE biology (such as this one), the sequences identified by RepeatMasker become the subject of downstream analysis.

Our pipeline was completed as follows: 1) Raw trimmed reads were queried using RepeatMasker against both RepBase and a custom repeat library, which contained known TEs from six other salamanders from the family Plethodontidae (*Aneides flavipunctatus*, *Batrachoseps nigriventris*, *Bolitoglossa occidentalis*, *Bolitoglossa rostrata*, *Desmognathus ochrophaeus*, and *Eurycea tynerensis*) as well as the hellbender salamander, *Cryptobranchus alleganiensis* from the family Cryptobranchidae (Sun, Arriaza, et al. 2012; Sun, Shepard, et al. 2012). Raw trimmed reads were also run through DnaPipeTE. 2) Repetitive sequences identified using DnaPipeTE were queried using RepeatMasker against the salamander library for annotation, as they are only annotated down to TE order level by DnaPipeTE itself. 3) A custom Perl script was used to parse out each RepeatMasker TE based on its base pair location within each read, as many reads contained multiple TEs. 4) Finally, the repetitive sequences detected by DnaPipeTE and TEs detected by RepeatMasker were combined for each species to characterize the total TE landscape for each species. Sequences that were identified as being repetitive, but not able to be classified, were referred to as “unknown repeats.” We calculated the percentage of each genome occupied by each TE superfamily, as well as by unknown repeats, by dividing the base pairs annotated to each superfamily by the total base pairs sequenced for each genome. We are assuming that the sequence data is a random subsample of the total genome sequence.

### Measuring Diversity of the Genomic TE Communities

TE diversity was measured for each species using both the Simpson’s and Shannon diversity indices in two different ways. In both methods, TE superfamilies are considered as species. In the first method, the total numbers of detected TE sequences annotated to each superfamily were considered as the number of individuals per “species.” In the second method, the total numbers of base pairs for each annotated superfamily were used for total presence of individuals per “species.” The second method differs from the first in that using base pair measurements takes into account the different sizes of TEs, as some can be significantly longer than others and therefore take up more space in the genome. Unknown repeats were excluded from the analysis, as were TEs that could only be annotated down to the level of Class (i.e. LTR). Simpson’s diversity index is expressed as the variable D, calculated by: 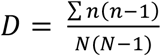 (Simpson 1949). D is the probability that two individuals at random pulled from a community will be from the same species. Since diversity decreases as D increases, this number is often expressed as 1 – D, or the Gini-Simpson’s index instead, which is more intuitive. The Shannon’s diversity index is represented by the variable H, which is calculated by: 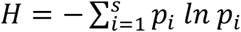 (Shannon 1948). The higher the value of H, the greater the diversity. Shannon’s diversity index is more sensitive to sample size and rarer species than is Simpson’s index (Mouillot and Leprêtre 1999), so the Shannon index may be a more accurate representation of genome diversity because of the presence of many low frequency repeats. However, with low-coverage data, rare repeats may go undetected, so we used both indices.

## RESULTS AND DISCUSSION

### Transposable Element Levels Are Similar Across Genome Sizes

For *Plethodon cinereus* and *P. glutinosus*, the MinION generated 4.15 Gb of data and 1.22 million reads, with an N50 of 6.59 kb. For *Plethodon vehiculum* and *P. idahoensis*, the MinION generated 2.11 Gb of data and 512,830 reads, with an N50 of 7.49 kb. These values are lower than expected based on MinION technology specs, but low data yield in applications of MinION sequencing to amphibian samples have also been reported in other studies (Menegon, et al. 2017; Pomerantz, et al. 2018; Lamichhaney, et al. 2021).

The combined outputs for RepeatMasker and DnaPipeTE identified the following numbers of repeats for each species: 2,153,518 for *P. cinereus*; 1,476,209 for *P. glutinosus*; 807,344 for *P. vehiculum*; and 898,214 for *P. idahoensis*. Between one and 94 individual TE sequences were annotated within single reads. In all four species, ~99% of the repeats were detected by RepeatMasker, and the remaining ~1% by DnaPipeTE.

Overall, the percentage of the genome composed of TEs ranged from 27% in *P. cinereus* to 32% in *P. idahoensis*, with an additional 6% - 8% composed of unknown repeats (Table 1). For each of the two genome expansion events encompassed by the pairwise comparisons — the lineage leading to *P. glutinosus* in the eastern clade and the lineage leading to *P. idahoensis* in the western clade — the percentage of the genome composed of recognizable TEs does not increase nearly as much as the genome size itself. The *P. glutinosus* genome is ~33% larger than that of *P. cinereus*, but the percentage of recognizable TEs is only 1% higher. Similarly, the *P. idahoensis* genome is ~40% larger than the *P. vehiculum* genome, but the percentage of recognizable TEs is only 2% higher. This result suggests that the increase in genome size is attributable to the accumulation of TEs that have persisted long enough to accumulate mutations and become unrecognizable, which in turn suggests decreased rates of TE deletion rather than recent bursts of TE proliferation. Interestingly, earlier DNA reassociation kinetic studies (i.e. Cot-curve comparisons) suggested that the percentage of repetitive DNA was much higher in the larger genome of *P. vehiculum* (80%) than in the smaller genome of *P. cinereus* (60%), a pattern that our results do not corroborate (Mizuno and Macgregor 1974).

**Table 1.** Total number of repeat sequences detected in each genome, and the percentage of the overall genome occupied by classifiable and unclassifiable repeats.

|  | **Genome Size (Gb)** | **Clade** | **Total # detected repeats** | **% genome occupied by classified TEs** | **% genome occupied by unknown repeats** |
| --- | --- | --- | --- | --- | --- |
| *P. cinereus* | 30.5 | Eastern | 2,153,518 | 27% | 7% |
| *P. glutinosus* | 40.4 | Eastern | 1,476,209 | 28% | 8% |
| *P. vehiculum* | 50.5 | Western | 807,344 | 30% | 7% |
| *P. idahoensis* | 70.6 | Western | 898,214 | 32% | 6% |

### Transposable Element Landscapes Are Similar Across Genome Sizes

All four species contained at least 52 TE superfamilies, which varied in relative abundance by 5 orders of magnitude within each genome (Table 2). Using both methods of calculating relative abundance — the total number of individual TE sequences and the total number of base pairs occupied by the TE superfamily — *Gypsy* (order LTR) was the most abundant in all four genomes, followed by L2 (order LINE) and DIRS (order DIRS). *Gypsy* accounted for 20% – 30% of the total repeats, and 27% – 33% of the total repeat base pairs; L2 accounted for 16% – 19% of the total repeats (15% – 19% base pairs); and DIRS accounted for 8% – 9% of the total repeats (11% – 13% base pairs) (Table 2). The least abundant superfamily across all four species was the DNA transposon Sola (TIR order), accounting for < 0.0007% of the total repeats/repeat base pairs. Overall, the most abundant TE superfamilies were dominated by retrotransposons; only PIF-Harbinger and Helitron exceeded 1% of the repeats in all four species. Unknown repeats accounted for 22% – 31% of the total repeats (15% - 22% base pairs).

**Table 2.** Percentage of total repetitive sequence in each genome that is composed of each TE superfamily as well as unknown repeats

| Order | Superfamily | % of total repeats (individual repeats) |  |  |  | % of total repeats (basepairs occupied by repeats) |  |  |  |
| --- | --- | --- | --- | --- | --- | --- | --- | --- | --- |
|  |  | *P. glutinosus* | *P. cinereus* | *P. idahoensis* | *P. vehiculum* | *P. glutinosus* | *P. cinereus* | *P. idahoensis* | *P. vehiculum* |
| Class I - Retrotransposon - Autonomous |  |  |  |  |  |  |  |  |  |
| LTR | *ERV* | 4.609 | 4.331 | 3.249 | 3.801 | 6.802 | 6.642 | 4.422 | 5.713 |
|  | *Gypsy* | 23.411 | 20.004 | 30.806 | 24.438 | 33.025 | 27.487 | 36.072 | 29.924 |
|  | *Bel-Pao* | 0.006 | 0.008 | 0.008 | 0.006 | 0.002 | 0.002 | 0.003 | 0.002 |
|  | *Copia* | 0.122 | 0.108 | 0.077 | 0.205 | 0.103 | 0.102 | 0.081 | 0.252 |
|  | *Bhikari* | 0.108 | 0.116 | 0.054 | 0.085 | 0.093 | 0.092 | 0.039 | 0.057 |
|  | *Unknown LTR* | 0.024 | 0.012 | 0.030 | 0.008 | 0.013 | 0.006 | 0.040 | 0.004 |
| DIRS | *DIRS* | 7.818 | 8.523 | 8.573 | 7.974 | 10.610 | 12.531 | 10.766 | 11.459 |
| LINE | *Ngaro* | 0.335 | 2.188 | 0.069 | 0.385 | 0.396 | 2.552 | 1.431 | 0.497 |
|  | *Penelope* | 1.965 | 1.242 | 0.710 | 0.970 | 0.981 | 0.849 | 0.484 | 0.628 |
|  | *Jockey* | 0.022 | 0.030 | 0.036 | 0.062 | 0.020 | 0.025 | 0.028 | 0.051 |
|  | *L1* | 2.870 | 3.287 | 4.435 | 5.473 | 1.575 | 1.905 | 2.607 | 3.137 |
|  | *L2* | 17.284 | 18.804 | 16.297 | 17.622 | 15.257 | 17.697 | 18.921 | 17.506 |
|  | *RTE* | 1.130 | 1.244 | 0.961 | 1.113 | 0.805 | 0.961 | 0.737 | 0.864 |
|  | *R1* | 0.002 | 0.001 | 0.015 | 0.000 | 0.001 | 0.001 | 0.013 | 0.000 |
|  | *R2* | 0.001 | 0.002 | 0.003 | 0.002 | 0.000* | 0.001 | 0.002 | 0.001 |
|  | *I* | 0.019 | 0.024 | 0.028 | 0.013 | 0.010 | 0.014 | 0.016 | 0.007 |
|  | *CR1* | 0.986 | 1.320 | 0.743 | 0.900 | 0.649 | 0.934 | 0.517 | 0.681 |
|  | *Proto2* | 0.000* | 0.000* | 0.000 | 0.000 | 0.000* | 0.000* | 0.000 | 0.000 |
|  | *Tad1* | 0.000 | 0.000 | 0.001 | 0.000 | 0.000 | 0.000 | 0.001 | 0.000 |
|  | *Unknown LINE* | 0.077 | 0.085 | 0.069 | 0.067 | 0.056 | 0.071 | 0.053 | 0.054 |
| Class I - Retrotransposon - Non-autonomous |  |  |  |  |  |  |  |  |  |
| SINE | *7SL* | 0.002 | 0.003 | 0.004 | 0.000 | 0.001 | 0.002 | 0.002 | 0.000 |
|  | *5S* | 0.027 | 0.027 | 0.125 | 0.131 | 0.009 | 0.010 | 0.046 | 0.054 |
|  | *tRNA* | 0.143 | 0.178 | 0.140 | 0.123 | 0.041 | 0.054 | 0.055 | 0.038 |
|  | *B4* | 0.007 | 0.012 | 0.005 | 0.006 | 0.002 | 0.004 | 0.001 | 0.002 |
|  | *Deu* | 0.897 | 1.040 | 1.151 | 1.829 | 0.478 | 0.580 | 0.624 | 1.003 |
|  | *MIR* | 0.368 | 0.428 | 0.120 | 0.271 | 0.176 | 0.217 | 0.060 | 0.145 |
|  | *Unknown SINE* | 0.001 | 0.001 | 0.002 | 0.001 | 0.001 | 0.001 | 0.002 | 0.001 |
| Class II - DNA Transposon - Subclass 1 |  |  |  |  |  |  |  |  |  |
| TIR | *hAT* | 1.588 | 1.258 | 0.384 | 0.421 | 1.325 | 0.877 | 0.209 | 0.235 |
|  | *Tc1-Mariner* | 0.629 | 0.909 | 1.305 | 0.688 | 0.382 | 0.625 | 0.828 | 0.426 |
|  | *PIF-Harbinger* | 1.511 | 1.582 | 2.379 | 2.302 | 1.261 | 1.287 | 1.452 | 1.868 |
|  | *PiggyBac* | 0.186 | 0.209 | 2.287 | 1.779 | 0.108 | 0.127 | 1.581 | 1.014 |
|  | *Sola* | 0.000* | 0.000* | 0.000* | 0.000* | 0.000* | 0.000* | 0.000* | 0.000* |
|  | *MuDR* | 0.014 | 0.016 | 0.024 | 0.042 | 0.008 | 0.009 | 0.013 | 0.048 |
|  | *P* | 0.006 | 0.004 | 0.002 | 0.004 | 0.002 | 0.002 | 0.001 | 0.002 |
|  | *Zisupton* | 0.058 | 0.075 | 0.125 | 0.058 | 0.029 | 0.046 | 0.051 | 0.030 |
|  | *Kolobok* | 0.019 | 0.029 | 0.012 | 0.006 | 0.011 | 0.019 | 0.005 | 0.003 |
|  | *Academ* | 0.015 | 0.020 | 0.004 | 0.009 | 0.011 | 0.015 | 0.003 | 0.007 |
|  | *Dada* | 0.006 | 0.011 | 0.005 | 0.004 | 0.003 | 0.006 | 0.002 | 0.002 |
|  | *Ginger* | 0.016 | 0.038 | 0.017 | 0.040 | 0.007 | 0.016 | 0.008 | 0.017 |
|  | *IS3EU* | 0.003 | 0.004 | 0.004 | 0.003 | 0.002 | 0.003 | 0.001 | 0.002 |
|  | *MULE* | 0.008 | 0.011 | 0.008 | 0.005 | 0.003 | 0.005 | 0.003 | 0.002 |
|  | *Merlin* | 0.004 | 0.004 | 0.006 | 0.003 | 0.001 | 0.001 | 0.002 | 0.001 |
|  | *CMC/En-Spm* | 0.099 | 0.159 | 0.219 | 0.171 | 0.046 | 0.076 | 0.097 | 0.014 |
|  | *Novosib* | 0.000* | 0.001 | 0.001 | 0.000* | 0.000* | 0.000* | 0.000* | 0.000* |
|  | *Zator* | 0.011 | 0.015 | 0.000 | 0.000 | 0.014 | 0.023 | 0.000 | 0.000 |
| Crypton | *Crypton* | 0.018 | 0.017 | 0.011 | 0.007 | 0.006 | 0.008 | 0.004 | 0.004 |
| Maverick | *Maverick* | 0.480 | 0.610 | 0.584 | 0.934 | 0.690 | 1.064 | 0.792 | 1.225 |
| Helitron | *Helitron* | 1.546 | 1.330 | 1.755 | 1.662 | 2.739 | 2.355 | 3.236 | 3.317 |
| Unknown Superfamilies |  | 0.103 | 0.108 | 0.117 | 0.122 | 0.071 | 0.084 | 0.077 | 0.097 |
| Unable to be classified |  | 31.337 | 30.385 | 21.898 | 26.200 | 22.142 | 20.545 | 14.574 | 19.586 |

\* indicates that the superfamily was detected at <0.001%

The percentage of the total genome occupied by each of the top 20 most abundant TE superfamilies is summarized in Figures 1 and 2. *Gypsy* accounted for 9.4% – 13.6% of the total genomic sequence in each genome. All four species had the same six most abundant TE superfamilies, in the same rank order: *Gypsy*, *L2*, *DIRS*, *ERV*, *Helitron*, and *L1*. Thus, we infer that, in both cases of genome expansion — on the lineage leading to *P. idahoensis* in the western clade, and on the lineage leading to *P. glutinosus* in the eastern clade — the most abundant superfamilies all contributed to genome expansion through an increase in copy number, reflecting increased proliferation and/or decreased deletion. There are more differences in rank abundance among the lower-frequency superfamilies, but with our low-coverage dataset, there is more error associated with those estimates.

**Figure 1.**
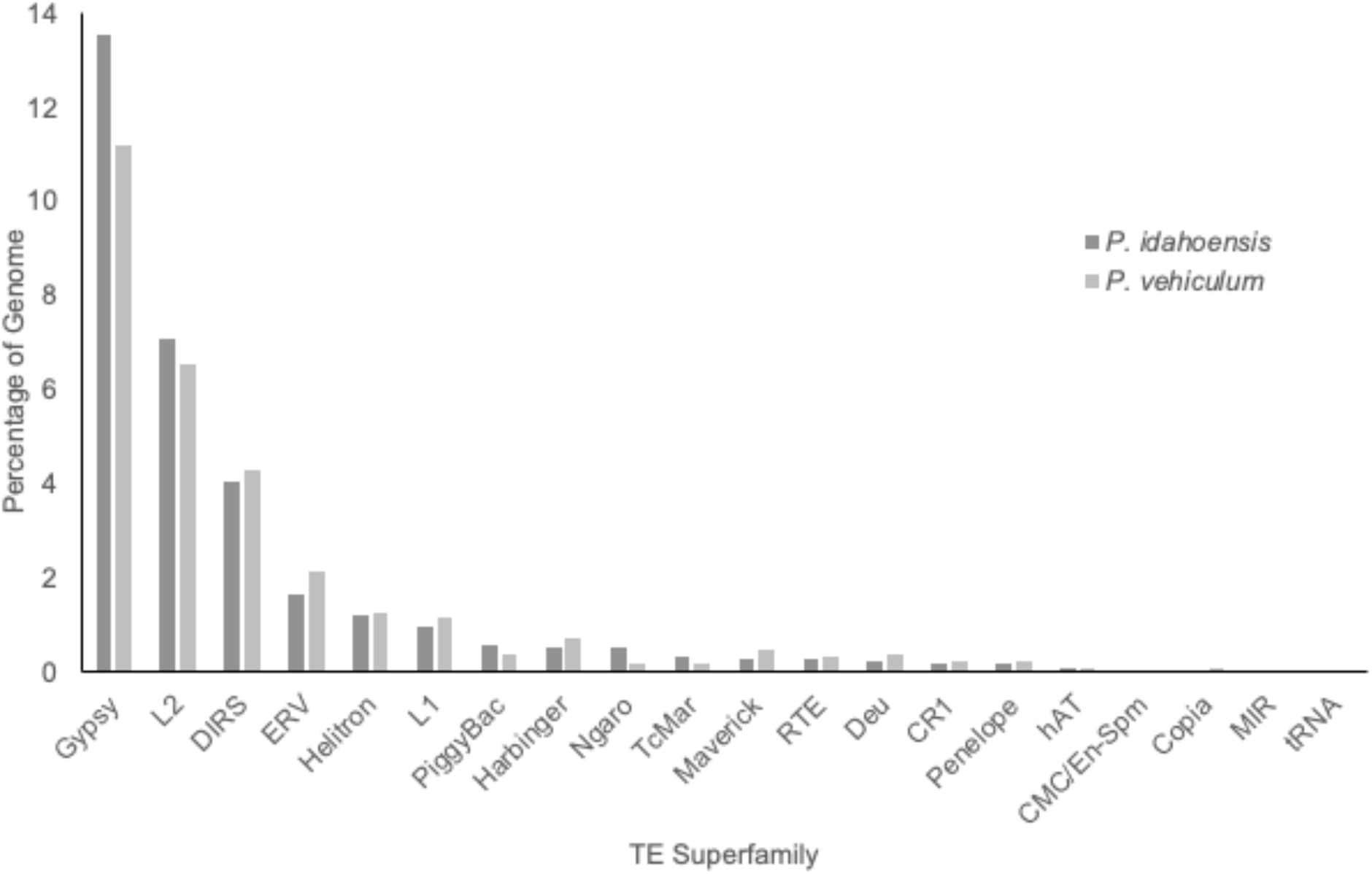
Percentage of the genome composed of the 20 most abundant TE superfamilies in *Plethodon idahoensis* and *Plethodon vehiculum* (western subclade). *P. idahoensis* has a larger genome than *P. vehiculum* (67.0 Gb vs. 46.4 Gb).

**Figure 2.**
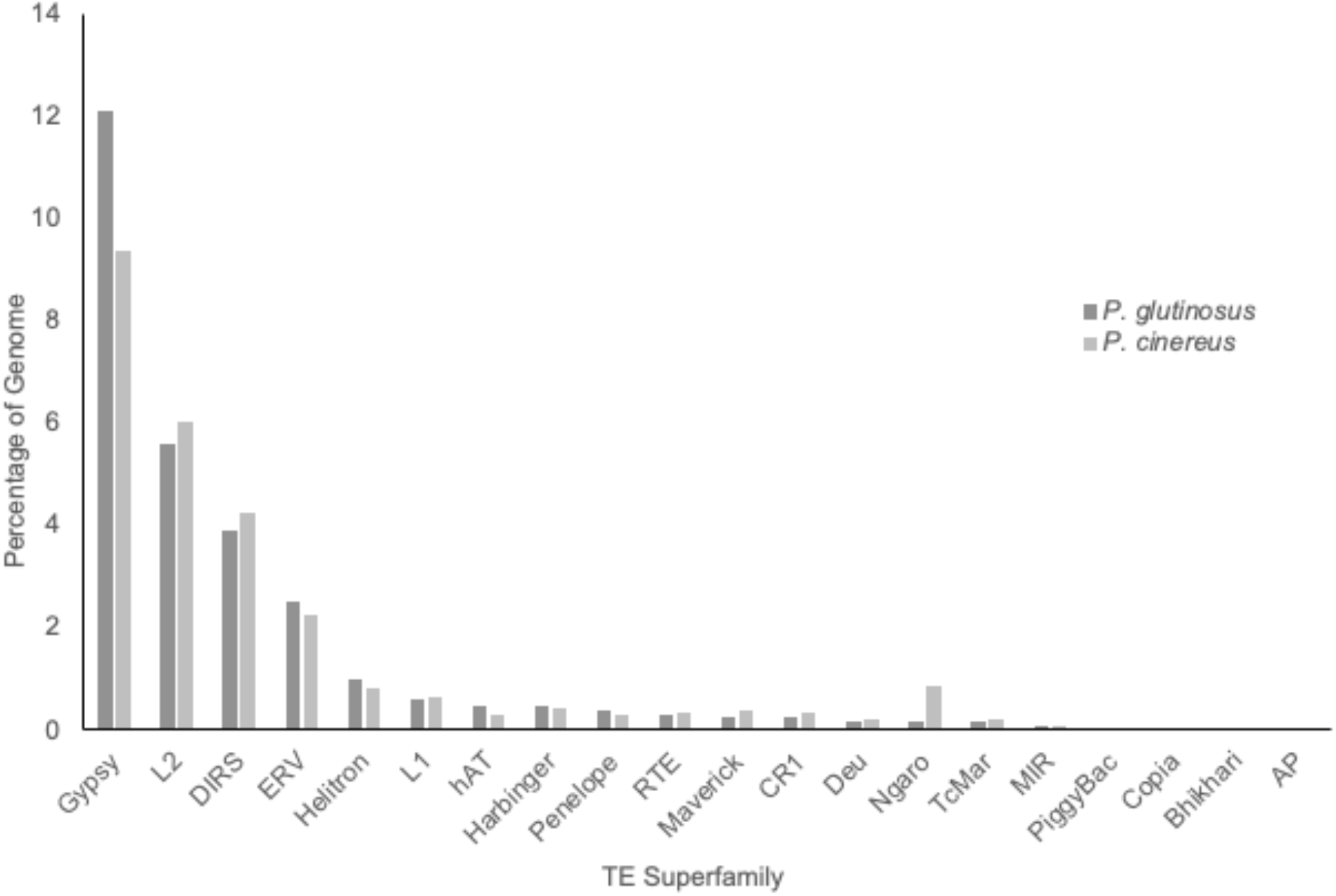
Percentage of the genome composed of the 20 most abundant TE superfamilies in *Plethodon glutinosus* and *Plethodon cinereus* (eastern subclade). *P. glutinosus* has a larger genome that *P. cinereus* (38.9 Gb vs. 29.3 Gb).

Overall, the four species contained nearly identical detected TE superfamilies. The two species from the eastern clade (*P. glutinosus* and *P. cinereus*) contained two superfamilies that were not detected in the western clade: Zator (a TIR DNA transposon) and Proto2 (a LINE retrotransposon). *Plethodon idahoensis* contained a superfamily that was not detected in the other three species, Tad1 (a LINE retrotransposon). *Plethodon vehiculum* was the only species in which the superfamily R1 (a LINE retrotransposon) was not detected. Because all of these are present at low relative abundances in the genomes, it is possible that their patterns of presence/absence across species reflect our inability to detect them with low-coverage data.

### Transposable Element Superfamily Diversity Remains Unchanged as Genome Size Increases in Salamanders

For both pairwise comparisons — *P. cinereus* and *P. glutinosus* in the eastern clade, and *P. vehiculum* and *P. idahoensis* in the western clade — the diversity indices were similar between the smaller and larger genome, demonstrating that a 10-Gb increase in genome size was not associated with a substantial change in TE community diversity measured at the superfamily level (Table 3). When the abundance of each TE superfamily was measured using TE copy number, the differences in Gini-Simpson’s index were 0.02 for the eastern clade (1-D = 0.82 and0.80 for *P. cinereus* and *P. glutinosus*, respectively) and 0.03 for the western clade (1-D = 0.81 and 0.78 for *P. vehiculum* and *P. idahoensis*, respectively). Using total base pairs occupied by each TE superfamily, the differences for both pairwise comparisons were 0.03. Using TE copy number, the differences in Shannon index were 0.11 for the eastern clade (H = 2.16 and 2.05 for *P. cinereus* and *P. glutinosus*, respectively) and 0.09 for the western clade (H = 2.12 and 2.03 for *P. vehiculum* and *P. idahoensis*, respectively). Using total base pairs occupied by each TE superfamily, the differences for both pairwise comparisons were 0.11. For context, a comparative study of TE superfamily diversity across vertebrates that encompassed species with smaller genomes found that the pufferfish *Takifugu rubripes* (0.4 Gb genome) had a Gini-Simpson index of 1.0 and Shannon index of 2.1, whereas the chicken *Gallus gallus* (1.3 Gb genome) had a Gini-Simpson index of 0.5 and Shannon index of 0.9, differing by 0.5 and 1.2, respectively (Wang, et al. 2021). These differences are an order of magnitude greater than the differences we report in salamanders. Other pairwise comparisons of TE superfamily diversity in vertebrate genomes that differ in relative size by about the same amounts as the salamanders we study here reveal both similar and different levels of diversity; for example, *G. gallus* (1.3 Gb) versus the frog *Xenopus tropicalis* (1.7 Gb) differ by 0.4 in Gini-Simpson index and 1.34 in Shannon index, whereas *X. tropicalis* versus the lizard *Anolis carolinensis* (2.2 Gb) differ by 0.01 in Gini-Simpson index and 0.17 in Shannon index (Wang, et al. 2021).

**Table 3.** Simpson and Shannon’s diversity indices for TE superfamily diversity

|  | Gini-Simpson Index (1-D)<br>Using total TE copy number | Shannon's Index (H)<br>Using total TE copy number | Gini-Simpson Index (1-D)<br>Using total base pair number | Shannon's Index (H)<br>Using total base pair number |
| --- | --- | --- | --- | --- |
| *Plethodon glutinosus* – 41.4 Gb | 0.80 | 2.05 | 0.76 | 1.88 |
| *Plethodon cinereus* – 30.5 Gb | 0.82 | 2.16 | 0.79 | 1.99 |
| *Plethodon idahoensis* – 71.7 Gb | 0.78 | 2.03 | 0.75 | 1.85 |
| *Plethodon vehiculum* – 50.5 Gb | 0.81 | 2.12 | 0.78 | 1.96 |

The diversity index values we report for *Plethodon* fall within the range reported for five species of salamanders that represent three families (Ambystomatidae, Cryptobranchidae, and Plethodontidae), two different types of datasets (whole-genome assembly and low-coverage 454 genome skimming), and a range of genome sizes (15 – 55 Gb) (Wang, et al. 2021). In that study, there was no correlation between genome size and TE superfamily diversity in salamanders. However, the species analyzed (*Desmognathus ochrophaeus*, *Batrachoseps nigriventris*, *Ambystoma mexicanum*, *Aneides flavipunctatus*, and *Cryptobranchus alleganiensis*) were phylogenetically quite divergent, including spanning the basal-most split in the salamander clade, and these large evolutionary distances could be associated with overall differences in genome biology that would obscure changes in TE diversity stemming from genome size. In addition, the deep evolutionary history encompassed by those five species captured increases and decreases along the lineages leading to the focal taxa (Sessions 2008). In contrast, our study system consisted of four more closely related species within the genus *Plethodon*, which are expected to have much more similar genomes overall. In addition, our taxon sampling yielded two pairwise comparisons in which the larger of the two genomes resulted from an increase in genome size (Newman, et al. 2016). Thus, the current study is a more powerful system for detecting decreases in TE diversity with increases in genome size. The fact that we do not see this pattern suggests that TE superfamily diversity remains high in enormous genomes. In addition, large genomes contain high levels of inactive and degraded TEs (Novák, et al. 2020), which are diverse in sequence. Thus, large genomes do not appear to be characterized by a low-diversity sequence community overall.

Our results suggest that the models that predict a decrease in diversity as genomes expand do not accurately capture the dynamics of TEs and their hosts in all cases. The richness of TE superfamilies may reach a maximum after the genome reaches a certain size (Elliott and Gregory 2015a) — we see 52 superfamilies represented in each *Plethodon* genome —and TE dynamics in large genomes may keep these superfamilies at the same evenness. Some of the suggested mechanisms predicting decreased diversity include competition among TEs to exploit host enzymes (Furano, et al. 2004) or evade host silencing machinery (Boissinot and Sookdeo 2016); our results suggest that these competitive interactions may not be relevant among TE superfamilies in large genomes. Finally, it is also possible that annotating only down to the superfamily level — considering every superfamily member as the same “species” — is not sensitive enough to detect relevant changes in TE diversity because each superfamily consists of multiple divergent families. For example, in mammals, one L1 family evades host silencing to be active at a time, whereas in lizards and other non-mammalian vertebrates, multiple active L1 families coexist, demonstrating differences in active TE diversity within the same superfamily (Boissinot and Sookdeo 2016). Overall, our results demonstrate that substantial increases in genome size occur without associated changes in TE diversity at the superfamily level.

## Declarations

### Funding

Funding was provided by the National Science Foundation (1911585 to RLM) and Colorado State University

### Conflicts of interest/Competing interests

Not applicable

### Availability of data and material

Sequence data is deposited in the NCBI Sequence Read Archive (SRA) under accession number PRJNA749318

### Code availability

Not applicable

### Ethics approval

Animal use was done in accordance with Colorado State University’s IACUC protocol number 17-7189A

### Consent to participate

NA

### Consent for publication

NA

## Acknowledgements

We acknowledge members of A. Haley’s Master’s Degree committee, D. Sloan and M. Stenglein, for valuable assistance. We thank members of the Mueller and Sloan labs for discussion. We thank M. Itgen for *Plethodon* tissues. Funding was provided by the National Science Foundation (1911585 to RLM) and Colorado State University.

## Notes

### Competing Interest Statement

The authors have declared no competing interest.

